# Sofosbuvir Terminated RNA is More Resistant to SARS-CoV-2 Proofreader than RNA Terminated by Remdesivir

**DOI:** 10.1101/2020.08.07.242156

**Authors:** Steffen Jockusch, Chuanjuan Tao, Xiaoxu Li, Minchen Chien, Shiv Kumar, Irina Morozova, Sergey Kalachikov, James J. Russo, Jingyue Ju

**Author notes:** SJ and CT contributed equally to this work.

## Abstract

SARS-CoV-2 is responsible for COVID-19, resulting in the largest pandemic in over a hundred years. After examining the molecular structures and activities of hepatitis C viral inhibitors and comparing hepatitis C virus and coronavirus replication, we previously postulated that the FDA-approved hepatitis C drug EPCLUSA (Sofosbuvir/Velpatasvir) might inhibit SARS-CoV-2.^1^ We subsequently demonstrated that Sofosbuvir triphosphate is incorporated by the relatively low fidelity SARS-CoV and SARS-CoV-2 RNA-dependent RNA polymerases (RdRps), serving as an immediate polymerase reaction terminator, but not by a host-like high fidelity DNA polymerase.^2,3^ Other investigators have since demonstrated the ability of Sofosbuvir to inhibit SARS-CoV-2 replication in lung and brain cells;^4,5^ additionally, COVID-19 clinical trials with EPCLUSA^6^ and with Sofosbuvir plus Daclatasvir^7^ have been initiated in several countries. SARS-CoV-2 has an exonuclease-based proofreader to maintain the viral genome integrity.^8^ Any effective antiviral targeting the SARS-CoV-2 RdRp must display a certain level of resistance to this proofreading activity. We report here that Sofosbuvir terminated RNA resists removal by the exonuclease to a substantially higher extent than RNA terminated by Remdesivir, another drug being used as a COVID-19 therapeutic. These results offer a molecular basis supporting the current use of Sofosbuvir in combination with other drugs in COVID-19 clinical trials.

## Introduction

SARS-CoV-2, the virus responsible for the COVID-19 pandemic, is a member of the Orthocoronavirinae subfamily.^9^ Coronaviruses and hepatitis C virus (HCV) are both positive-sense single-strand RNA viruses,^10,11^ with comparable mechanisms requiring an RNA-dependent RNA polymerase (RdRp) for its genome replication and transcription. Potential inhibitors have been investigated to target various steps in the Coronavirus infectious cycle, including the viral replication machinery.^10^ However, as of now, no effective therapeutic is available to treat serious coronavirus infections such as COVID-19. The RdRp is one of the key targets for antiviral drug development. This RNA polymerase is highly conserved at the amino acid level in the active site among different positive sense RNA viruses, including coronaviruses and HCV.^12^ Viral RdRps are highly error-prone,^13^ and therefore have the ability to accept modified nucleotide analogues as substrates. Nucleotide and nucleoside analogues that inhibit polymerases comprise an important group of antiviral agents.^14-17^

In late January of 2020, before COVID-19 reached pandemic status, based on our analysis of the molecular structures and activities of hepatitis C viral inhibitors and a comparison of hepatitis C virus and coronavirus replication, we postulated that the FDA-approved hepatitis C drug EPCLUSA (Sofosbuvir/Velpatasvir) might inhibit SARS-CoV-2.^1^ Using a computational approach, Elfiky predicted that Sofosbuvir, IDX-184, Ribavirin, and Remdesivir might be potent drugs against COVID-19.^18^ We subsequently demonstrated that Sofosbuvir triphosphate is incorporated by the relatively low fidelity SARS-CoV and SARS-CoV-2 RdRps, serving as an immediate polymerase reaction terminator, but not by a host-like high fidelity DNA polymerase.^2,3^ We also reported that a library of additional nucleotide analogues with a variety of structural and chemical features terminate RNA synthesis catalyzed by polymerases of coronaviruses that cause SARS and COVID-19.^19^ Gordon et al. performed a kinetic study, including the determination of Km values for triphosphates of Remdesivir, Sofosbuvir and other nucleotide analogues.^20^ Jácome et al. recently recommended Sofosbuvir as a possible antiviral for COVID-19, based on structural studies and bioinformatic analysis.^21^ By comparing the RNA genomes of HCV and SARS-CoV-2, Buonaguro et al. suggested that Sofosbuvir might be an optimal nucleotide analogue to repurpose for COVID-19 treatment.^22^ Other investigators have since demonstrated the ability of Sofosbuvir to inhibit SARS-CoV-2 replication in lung and brain cells,^4,5^ and COVID-19 clinical trials with EPCLUSA^6^ and with Sofosbuvir plus Daclatasvir^7^ have been initiated in several countries. Recently, Sadeghi et al. reported encouraging results from a clinical trial using Sofosbuvir (SOF) and Daclatasvir (DCV) as a potential combination treatment for moderate or severe COVID-19 patients.^23^ In a study involving 66 patients, these investigators showed that SOF/DCV treatment increased 14-day clinical recovery rates and reduced the length of hospital stays. They indicated that larger well controlled randomized trials are necessary to confirm these results.

Unlike many other RNA viruses, SARS-CoV and SARS-CoV-2 have very large genomes (∼30 kb) that encode a 3’-5’ exonuclease (nsp14) that carries out proofreading;^24,25^ this activity is enhanced by the nsp10 cofactor.^8^ This exonuclease-based proofreader increases replication fidelity by removing mismatched nucleotides to maintain the viral genome integrity.^26^ Any effective antiviral targeting the SARS-CoV-2 RdRp must display a certain level of resistance to this proofreading activity. Remdesivir (RDV), a nucleotide analogue targeting the SARS-CoV-2 RdRp, is currently used for the treatment of COVID-19 under FDA emergency authorization.^27^ The active triphosphate form of Remdesivir possesses a ribose with an OH group at both the 2’ and 3’ positions, while Sofosbuvir triphosphate has a 2’-F,Me-deoxyribose. The chemical stability of deoxyribose is much higher than that of the ribose. Although we and others have demonstrated that Sofosbuvir triphosphate (SOF-TP) can terminate the reaction catalyzed by coronavirus RdRps, it is not known whether Sofosbuvir terminated RNA will offer any resistance to the SARS-CoV-2 exonuclease-based proofreader. We report here that Sofosbuvir terminated RNA resists removal by the exonuclease complex (nsp14/nsp10) to a substantially higher extent than RNA terminated by Remdesivir. We demonstrate for the first time that upon incorporation of the triphosphate form of Sofosbuvir into RNA by the SARS-CoV-2 RdRp, SOF is removed at a lower rate by the SARS-CoV-2 exonuclease complex, relative to RDV and UMP. Thus, not only does the active triphosphate of Sofosbuvir serve as an efficient terminator of the polymerase, but SOF terminated RNA confers a substantial level of resistance to excision by exonuclease. Thus, the current use of Sofosbuvir in combination with other drugs in COVID-19 clinical trials^6,7^ is supported by these molecular level results.

## Results and Discussion

Sofosbuvir (Fig. 1a), a pyrimidine nucleotide analogue prodrug, has a hydrophobic masked phosphate group that enhances its ability to enter host cells. The prodrug is subsequently converted into an active triphosphate form (SOF-TP; 2’-F,Me-UTP) by cellular enzymes, enabling it to inhibit the HCV RdRp NS5B.^28,29^ SOF-TP is incorporated into RNA by the viral RdRp, and due to the presence of fluoro and methyl modifications at the 2’ position, inhibits further RNA chain extension, which halts RNA replication and viral multiplication. A related purine nucleotide ProTide^30^ prodrug, Remdesivir (Fig. 1b), originally developed for the treatment of Ebola virus infections, though not successfully, is being used for COVID-19 treatment. Unlike Sofosbuvir (Fig. 1a), Remdesivir has both 2’- and 3’-OH groups (Fig. 1b), and a cyano group at the 1’ position which is responsible for RdRp inhibition.

**Fig. 1.**
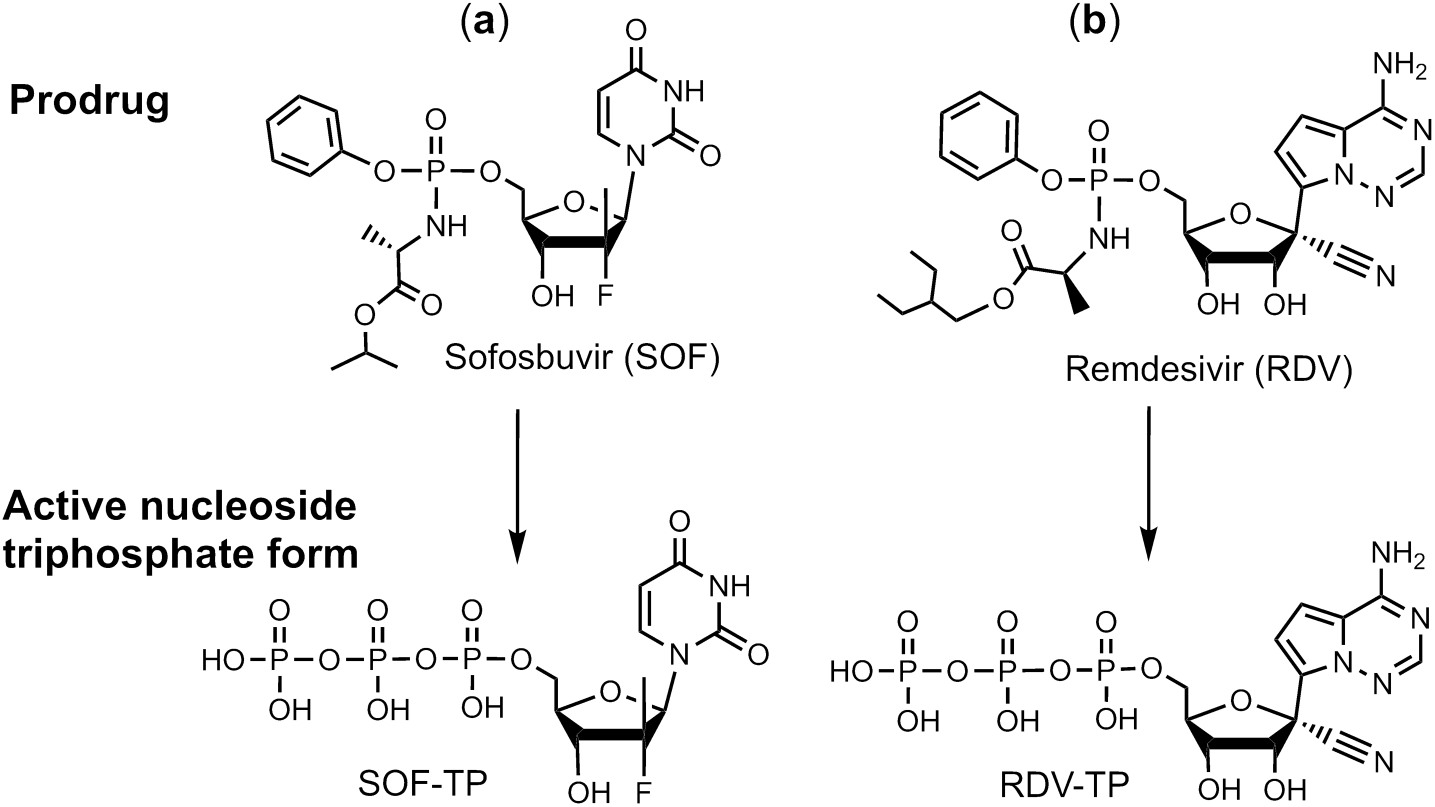
Comparison of structures of the prodrugs (a) Sofosbuvir (SOF) and (b) Remdesivir (RDV) and their active triphosphate forms (SOF-TP and RDV-TP). Top: Prodrug (phosphoramidate) forms; Bottom: Active triphosphate forms.

Analyzing the structures of the active triphosphate forms of Sofosbuvir (Fig. 1a) and Remdesivir (Fig. 1b), both of which have been shown to inhibit the replication of specific RNA viruses (Sofosbuvir for HCV, Remdesivir for SARS-CoV-2), we noted that the 2’-modifications in Sofosbuvir (the fluoro and methyl groups) are smaller than the 1’-cyano group and the 2’-OH group in Remdesivir. The bulky cyano group in close proximity to the 2’-OH may lead to steric hindrance, thereby impacting the polymerase reaction termination efficiency of the activated form of Remdesivir. It was recently reported that, using the MERS-CoV and SARS-CoV-2 RdRps, RDV-TP had higher incorporation efficiency than ATP. ^20,31^ However, RDV-TP does not cause immediate polymerase reaction termination; rather, it leads to delayed polymerase termination, likely due to its 1’-cyano group and free 2’-OH and 3’-OH groups.

To demonstrate whether the RNA terminated by the triphosphate forms of Sofosbuvir (SOF-TP) and Remdesivir (RDV-TP) have the potential to resist the SARS-CoV-2 proofreading activity, we carried out polymerase extension reactions followed by exonuclease digestion reactions. First, using the replication complex assembled from SARS-CoV-2 nsp12 (the viral RdRp) and nsp7 and nsp8 proteins (RdRp cofactors), the nucleotide analogues were incorporated at the 3’ end of the double-stranded segment of the RNA template-loop-primer shown at the top of Fig. 2. Because the template strand consists of an A at the next position, SOF-TP (a UTP analogue) or UTP are incorporated in a single-nucleotide extension reaction. On the other hand, because the next base in the template strand after the A is a U, in order to incorporate the ATP analogue RDV-TP, a two-nucleotide extension (UTP followed by RDV-TP) occurs. After purification to remove the RdRp and nucleoside triphosphates, the extended RNA is treated with the SARS-CoV-2 exonuclease complex consisting of SARS-CoV-2 nsp14 (the viral exonuclease) and nsp10 (an exonuclease cofactor) to determine whether excision of the incorporated nucleotide analogues takes place. Our prediction was that SOF would be at least partially protected from the exonuclease due to the presence of 2’-fluoro and 2’-methyl groups in place of the 2’-OH group, but that RDV and UMP, both of which have a 2’-OH and 3’-OH, would be less protected. This was indeed what we observed as described below.

**Fig. 2.**
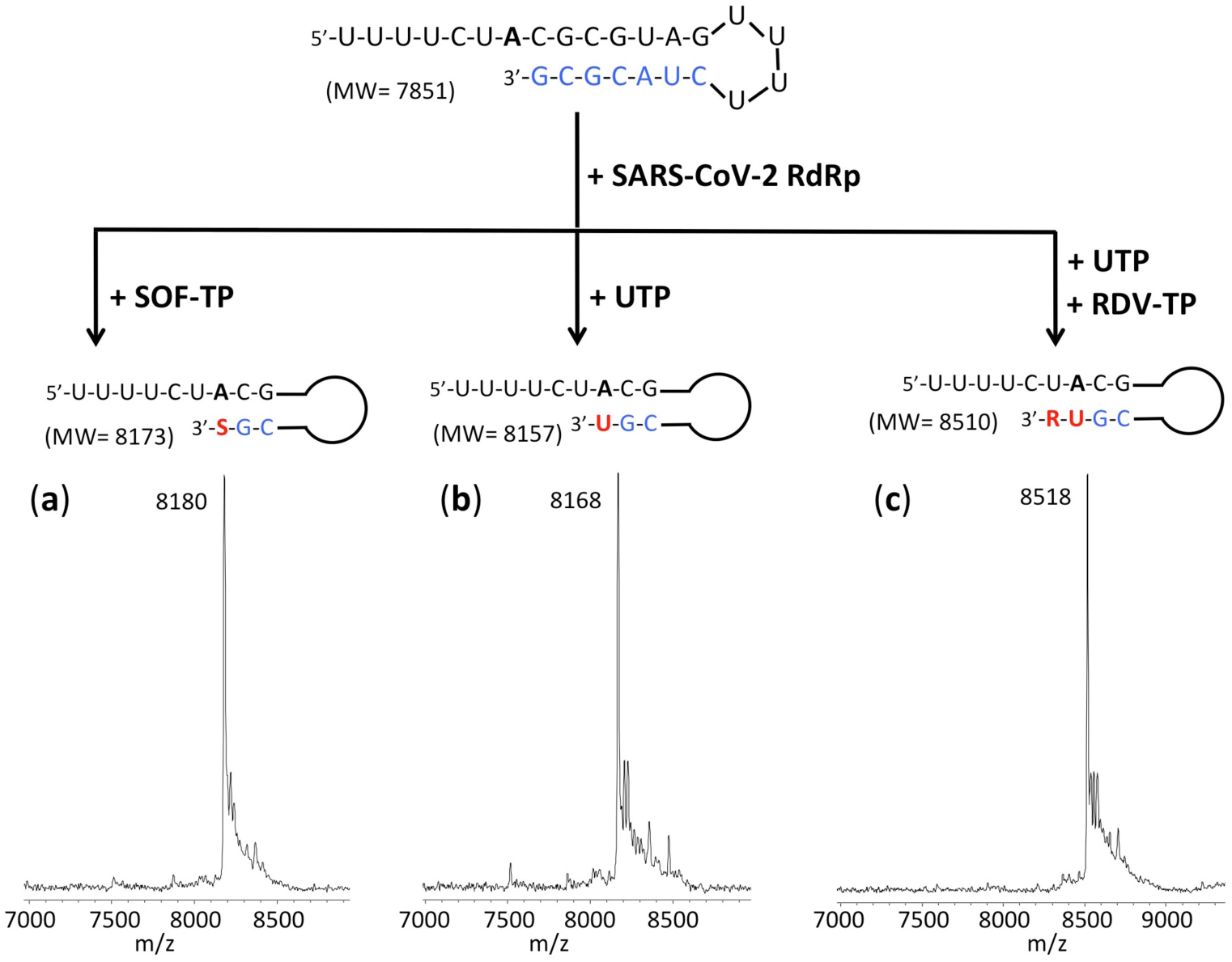
Polymerase extension reactions with SOF-TP, UTP or UTP+RDV-TP to produce RNA products. The sequence of the RNA template-loop-primer used for these polymerase extension reactions is shown at the top of the figure. Polymerase extension reactions were performed by incubating (a) SOF-TP, (b) UTP and (c) UTP + RDV-TP with pre-assembled SARS-CoV-2 polymerase (nsp12, nsp7 and nsp8) and the indicated RNA primer-loop-template, followed by detection of reaction products by MALDI-TOF MS. The accuracy for m/z determination is approximately ± 10 Da.

We performed polymerase extension reactions with SOF-TP, UTP, and RDV-TP + UTP, following the addition of the pre-annealed RNA template-loop-primer to a pre-assembled mixture of nsp12, nsp7 and nsp8. The extended RNA products from the reaction were subjected to MALDI-TOF-MS analysis to confirm that the expected RNA products were formed. The sequence of the RNA template-loop-primer used for the polymerase extension assay, which has previously been described,^32^ is shown at the top of Fig. 2.

As shown in Fig. 2a, following incubation with the SARS-CoV-2 replication complex, complete conversion of the RNA template-loop-primer (7851 Da expected) to the Sofosbuvir-terminated extension product was observed (8180 Da observed, 8173 Da expected). Similarly, as shown in Fig. 2b, quantitative extension was seen with UTP (8168 Da observed, 8157 Da expected) and as demonstrated in Fig. 2c, quantitative extension by UMP and RDV occurred (8518 Da observed, 8510 Da expected).

The above RNA extension products were purified and then incubated with the exonuclease complex (nsp14 and nsp10); the results are presented in Fig. 3. The MS trace for the Sofosbuvir extension product in the absence of exonuclease (0 min) is shown in Fig. 3a. Only the expected peak at 8183 Da (8173 Da expected) is observed. After 5 min treatment with the nsp14/nsp10 exonuclease complex (Fig. 3b), there is minimal appearance of cleavage products (e.g., the small 6558 Da peak representing cleavage of 5 nucleotides). Even after 30 min exonuclease treatment (Fig. 3c), there is a significant amount of the intact Sofosbuvir terminated RNA remaining, and the appearance of lower molecular weight peaks, for example at 6867 Da and 6561 Da (removal of 4 and 5 nucleotides from the 3’ end respectively). The MS result for the purified RNA extended with UMP is shown in Fig. 3d (8165 Da observed, 8157 expected). In addition, a smaller peak at 8472 represents mismatch incorporation of an additional U; this is likely due to the high concentration of UTP used and the relatively low fidelity of the RdRp.^13^ After 5 min treatment with the exonuclease complex (Fig. 3e), there is substantial cleavage as indicated by the peaks at 7211 Da, 6864 Da and 6559 Da (3, 4 and 5 nucleotide cleavage products, respectively). At 30 min (Fig. 3f), the extended RNA product peak completely disappears with the presence of only cleavage fragment peaks. Finally, the UMP + RDV extended RNA is shown in Fig. 3g with a single peak (8518 Da observed, 8510 Da expected). After 5 min exonuclease treatment (Fig. 3h), the RNA extension product peak is substantially reduced, with conversion to smaller fragments, e.g., 7210 Da, 6865 Da and 6559 Da (removal of 3, 4 or 5 nucleotides from the 3’ end). At 30 min (Fig. 3i), there is no visible RDV terminated RNA peak remaining, with only cleavage product peaks observed, e.g., at 7209 Da, 6863 Da, 6558 Da and 5923 Da (cleavage of 3, 4, 5 and 7 nucleotides respectively). Comparing the results in Fig. 3b with Fig. 3h, and Fig. 3c with Fig. 3i, it is clear that there is substantially more excision of Remdesivir than Sofosbuvir, with subsequent further cleavage to smaller RNA products. By comparing the results in Fig. 3d-i, it is also apparent that RDV is removed by the viral exonuclease more rapidly than UMP.

**Fig. 3.**
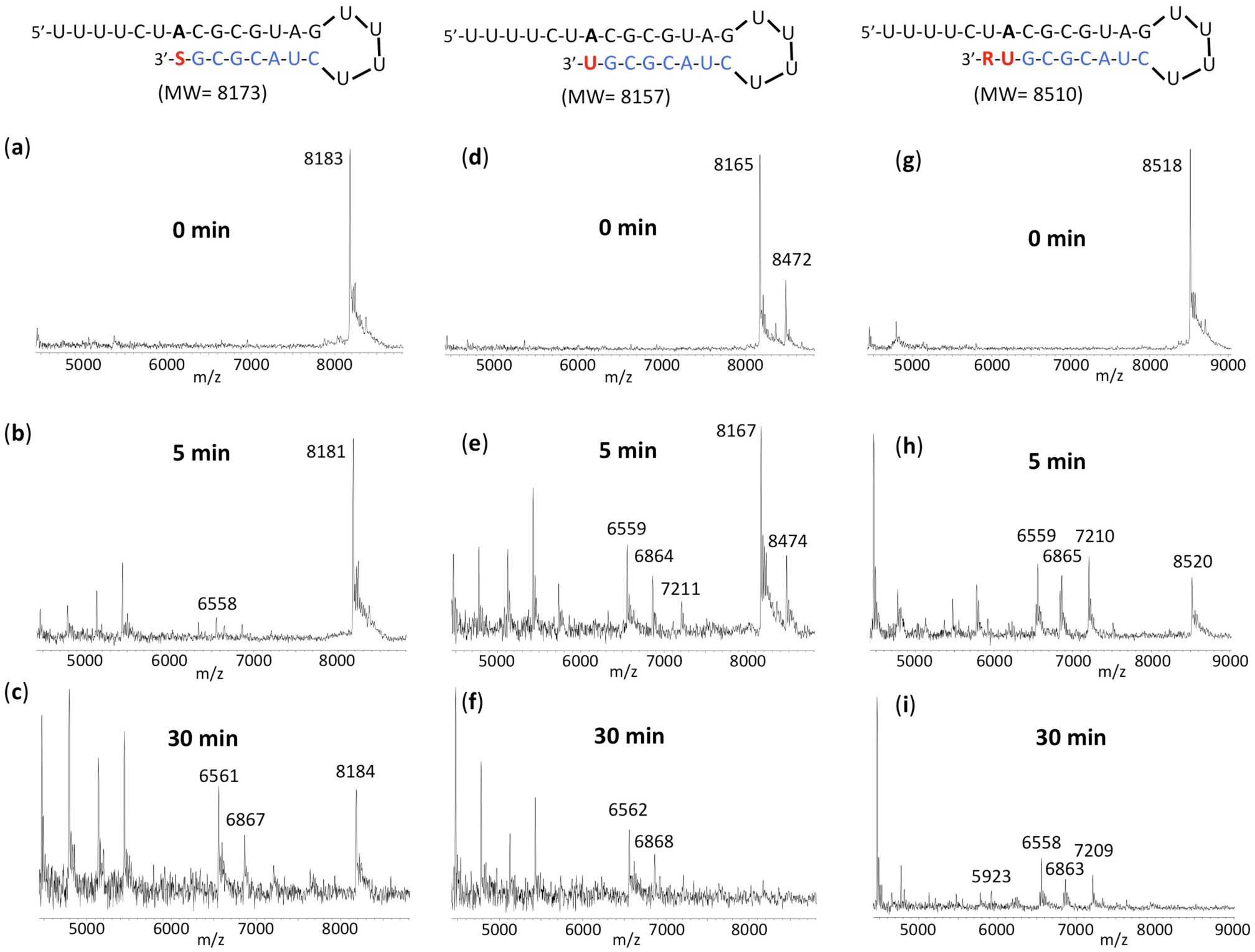
Treatment of the RNA products with exonuclease to determine relative excision of Sofosbuvir, UMP and Remdesivir. Untreated products (0 min) are shown in (a) for SOF extended RNA, (d) for UMP extended RNA and (g) for UMP plus RDV extended RNA. Exonuclease reactions were performed by incubating the purified RNA products, generated using the same procedure as in Fig. 2, with preassembled SARS-CoV-2 exonuclease complex (nsp14 and nsp10) for 5 min (b, e and h) or 30 min (c, f, i), followed by detection of reaction products by MALDI-TOF MS. The signal intensities were normalized to the highest peak within each time series. The accuracy for m/z determination is approximately ± 10 Da.

In order to further compare the relative excision among the different extended RNAs, in Fig. 4, similar experiments were carried out as in Fig. 3, but the exonuclease treated SOF and UMP extended RNA products (Fig. 4a-b) were combined for purification, followed by MALDI-TOF-MS analysis. The MS spectrum for the untreated RNA products is shown in Fig. 4a. In the inset it is possible to differentiate the RNA extended with UMP (8163 Da observed, 8157 Da expected) and SOF (8180 Da observed, 8173 Da expected). The MS spectrum for the 15 min exonuclease-treated RNA products is shown in Fig. 4b. It is clear in the inset that the RNA peak representing UMP extension (8164 Da) is reduced to a substantially greater extent than the peak representing the SOF extended RNA (8180 Da). In addition, peaks indicating cleavage by 3-7 nucleotides are present (7208 Da, 6862 Da, 6557 Da, 6227 Da and 5921 Da). Similarly, the exonuclease treated SOF and UMP+RDV extended RNA products (Fig. 4c-e) were combined for purification, followed by MALDI-TOF-MS analysis. The MS spectrum for the untreated RNA products is shown in Fig. 4c. The peak at 8179 Da represents the SOF extended product (8173 Da expected) and the peak at 8517 Da represents the UMP+RDV extended product (8510 Da expected). The MS spectrum for the 5 min exonuclease-treated RNA products is shown in Fig. 4d. There is some reduction in the height of the SOF-extended RNA peak, and a much more substantial reduction in the height of the UMP+RDV extended RNA peak. The peaks at 7209 Da, 6864 Da and 6559 Da represent exonuclease fragments after cleavage of 3-5 nucleotides, presumably mainly from the UMP+RDV extended RNA. In the case of the 30 min exonuclease-treated RNA products (Fig. 4e), there is still some SOF extended RNA remaining (8183 Da) while the UMP+RDV extended RNA peak is at baseline level. The peaks at 7209 Da, 6864 Da, 6559 Da, 6230 Da and 5923 Da represent exonuclease fragments after cleavage of 3-7 nucleotides. This experiment confirms the results in Fig. 3, that Sofosbuvir is more protected from cleavage by the SARS-CoV-2 exonuclease than UMP or Remdesivir.

**Fig. 4.**
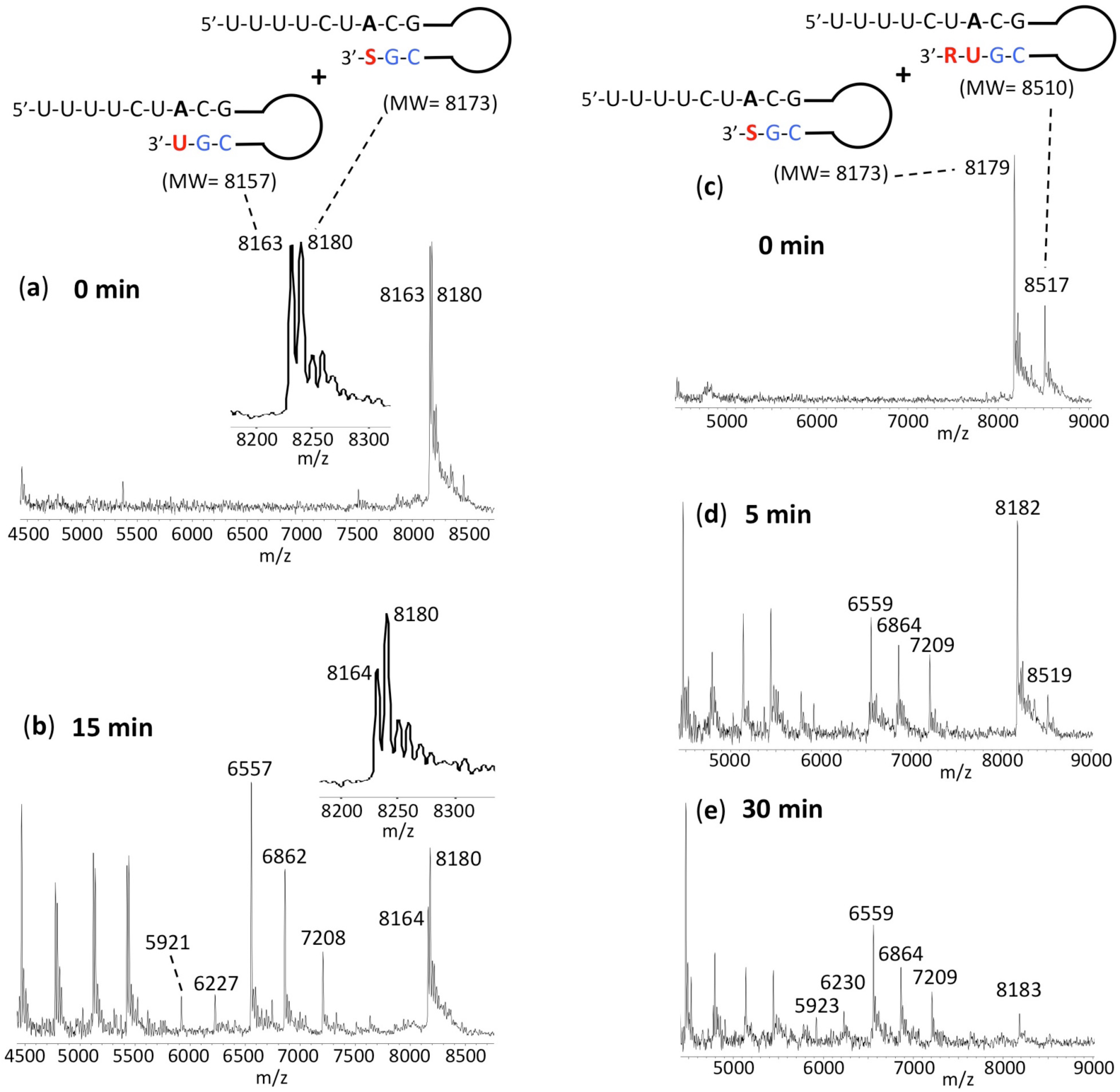
Treatment of the RNA products with exonuclease and then combined for analysis to determine relative excision of Sofosbuvir, UMP and Remdesivir. Untreated products (0 min) are shown in (a) for SOF and UMP extended RNA and (c) for SOF and UMP+RDV extended RNA. Exonuclease reactions were performed by incubating the purified RNA products with preassembled SARS-CoV-2 exonuclease complex (nsp14 and nsp10). The SOF and UMP extended RNAs were treated with the exonuclease complex for 15 min and then combined for purification followed by MALDI-TOF-MS (Fig. 4b). The insets in Fig. 4a and Fig. 4b are an enlargement of the 8200-8300 Da portion of the spectrum. The SOF and UMP+RDV extended RNAs were treated with the exonuclease complex for 5 min or 30 min and combined for purification and MALDI-TOF-MS (Fig. 4d and Fig. 4e, respectively). The signal intensities were normalized to the highest peak within each time series. The accuracy for m/z determination is approximately ± 10 Da.

## Conclusion

Our assay described above, which combines extension of RNA with the nucleotide analogue by the polymerase complex, and subsequent treatment with the exonuclease complex, can be used to test any nucleotide inhibitor that can be incorporated into SARS-CoV-2 RNA for its ability to resist exonuclease activity. The template-loop-primers can be designed to allow 1-step extension and termination by nucleotide analogues containing A, C, G or U bases. With MALDI-TOF MS, it is possible to distinguish extension and cleavage of nucleotides differing by ∼10 Da, especially when included in the same sample as in Fig. 4, a size difference that is not resolvable by assays that utilize gel electrophoresis.

The inhibition of SARS-CoV-2 in cell culture by Sofosbuvir has recently been reported. Sacramento et al. demonstrated that Sofosbuvir inhibited SARS-CoV-2 replication in human hepatoma-derived (Huh-2) and Type II pneumocyte-derived (Calu-3) cells with EC50 values of 6.2 and 9.5 µM, respectively.^4^ Mesci et al. showed that Sofosbuvir could protect human brain organoids from SARS-CoV-2 infection.^5^ Considering its low toxicity, its ability to be rapidly activated to the triphosphate form by cellular enzymes, and the high intracellular stability of this active molecule, Sayad et al. initiated a COVID-19 treatment clinical trial with Sofosbuvir and Velpatasvir, which together form the combination HCV drug EPCLUSA.^6^ Izzi et al. also advocated the use of Sofosbuvir and Velpatasvir for the treatment of COVID-19.^33^ The Sofosbuvir triphosphate has been shown to persist for over 24 h in hepatocytes.^34^ Remdesivir has been approved under FDA emergency use authorization,^27^ and is undergoing safety and efficacy trials for COVID-19; on the other hand, the FDA-approved hepatitis C drug Sofosbuvir is widely available and has a well-defined safety and clinical profile. Sofosbuvir and Velpatasvir form the HCV combination drug EPCLUSA. Velpatasvir inhibits NS5A, an essential protein required for HCV replication. NS5A enhances the function of RNA polymerase NS5B during viral RNA genome replication.^35,36^ The drug Daclatasvir also inhibits this protein.^37^ Daclatasvir was also shown to reduce SARS-CoV-2-induced enhancement of TNF-α and IL-6, key contributors to the cytokine storm, observed in some COVID-19 patients.^4^ Because Velpatasvir and Daclatasvir share very similar core structures and target the same NS5A protein in HCV, and Daclatasvir has also been shown to inhibit SARS-CoV-2 replication^4^ and is currently in COVID-19 clinical trial,^7^ it is plausible that Velpatasvir and other drugs in this class, such as Ledipasvir,^38^ Elbasvir,^39^ Ombitasvir^40^ and Pibrentasvir,^41^ will display similar inhibitory activity for SARS-CoV-2.

In summary, we demonstrated that RNA terminated with Sofosbuvir displays substantial resistance to excision by the SARS-CoV-2 exonuclease complex, relative to excision of the natural nucleotide UMP. Moreover, Sofosbuvir appears to show substantially more protection against SARS-CoV-2 exonuclease activity than Remdesivir. These results offer a molecular basis for the use of Sofosbuvir in combination with other drugs in clinical trials for prevention and treatment of COVID-19.

## Methods

The RdRp of SARS-CoV-2, referred to as nsp12, and its two protein cofactors, nsp7 and nsp8, shown to be required for the processive polymerase activity of nsp12, were cloned and purified as described.^3^ The 3’-exonuclease, referred to as nsp14, and its protein cofactor, nsp10, were purchased from LSBio (Seattle, WA). Sofosbuvir triphosphate (PSI-7409) was purchased from Sierra Bioresearch (Tucson, AZ). Remdesivir triphosphate (GS-443902) was purchased from MedChemExpress (Monmouth Junction, NJ). UTP was purchased from Fisher Scientific. The RNA oligonucleotide (template-loop-primer) was purchased from Dharmacon (Horizon Discovery, Lafayette, CO).

### Extension reactions with RNA-dependent RNA polymerase

The RNA template-loop-primer (sequence shown in Fig. 2) was annealed by heating to 75°C for 3 min and cooling to room temperature in 1x reaction buffer. The RNA polymerase mixture consisting of 11 µM nsp12, 66 µM nsp8 and 33 µM nsp7 was incubated for 15 min at room temperature in a 1:6:3 ratio in 1x reaction buffer. Then 10 µL of the annealed RNA template-loop-primer solution (5 µM) in 1x reaction buffer was added to 9 µL of the RNA polymerase mixture and incubated for an additional 10 min at room temperature. Finally, 1 µL of a solution containing 10 mM SOF-TP (Fig. 2a), 1 mM UTP (Fig. 2b), or 1 mM UTP + 1 mM RDV-TP (Fig. 2c) in 1x reaction buffer was added and incubation was carried out for 2 hr at 30°C. The final concentrations of reagents in the 20 µL extension reactions were 5 µM nsp12, 30 µM nsp8, 15 µM nsp7, 2.5 µM RNA template-loop-primer, 500 µM SOF-TP (Fig. 2a), 50 µM UTP (Fig. 2b) and 50 µM UTP + 50 µM RDV-TP (Fig. 2c). The 1x reaction buffer contains the following reagents: 10 mM Tris-HCl pH 8, 10 mM KCl, 2 mM MgCl2 and 1 mM β-mercaptoethanol. Desalting of the reaction mixture was performed with an Oligo Clean & Concentrator kit (Zymo Research) resulting in ∼10 µL purified aqueous RNA solutions. 2 µL of each solution were subjected to MALDI-TOF-MS (Bruker ultrafleXtreme) analysis, following a previously described method.^42^ The remaining ∼8 µL extended template-loop-primer solutions were used to test exonuclease activity as described below.

### Exonuclease reactions

The extended RNA template-loop-primers from above (sequences shown in Fig. 3) were annealed by heating to 75°C for 3 min and cooling to room temperature in 1x exonuclease reaction buffer. The exonuclease nsp14 (500 nM) and its protein cofactor, nsp10 (2 µM), were incubated for 15 min at room temperature in a 1:4 ratio in 1x exonuclease reaction buffer. Then 10 µL of the annealed extended RNA template-loop-primer solution (1 - 1.6 µM) in 1x exonuclease reaction buffer was added to 10 µL of the exonuclease mixture and incubated at 37°C for 5, 15 or 30 min. The final concentrations of reagents in the 20 µL reactions were 250 nM nsp14, 1 µM nsp10 and 500 - 800 nM extended RNA template-loop-primer. The 1x exonuclease reaction buffer contains the following reagents: 40 mM Tris-HCl pH 8, 1.5 mM MgCl2, 50 µM ZnCl2 and 5 mM DDT. Following desalting using an Oligo Clean & Concentrator (Zymo Research), the samples were subjected to MALDI-TOF-MS (Bruker ultrafleXtreme) analysis.

## Acknowledgements

This research is supported by Columbia University, a grant from the Jack Ma Foundation, a generous gift from the Columbia Engineering Member of the Board of Visitors Dr. Bing Zhao, and Fast Grants to J.J. We thank Dr. Robert Kirchdoerfer for constructing the RdRp (nsp12, nsp7 and nsp8).

## Author Contributions

J.J. conceived and directed the project; the approaches and assays were designed and conducted by J.J., S.J., C.T., X.L., M.C, S.K., I.M., Se.K. and J.J.R., and Data were analyzed by all authors. All authors wrote the manuscript.

